# Quaternary structural convergence and structural diversification of prion assemblies at the early replication stage

**DOI:** 10.1101/583781

**Authors:** Angélique Igel-Egalon, Florent Laferrière, Mohammed Moudjou, Mathieu Merzach, Tina Knäpple, Laetitia Herzog, Fabienne Reine, Marie Doumic, Human Rezaei, Vincent Béringue

## Abstract

Aggregation of misfolded forms from host-encoded proteins is key to the pathogenesis of a number of neurodegenerative disorders, including prion diseases, Alzheimer’s disease and Parkinson’s disease. In prion diseases, the cellular prion protein PrP^C^ can misfold into PrP^Sc^ and auto-organize into conformationally distinct assemblies or strains. A plethora of observations reports the existence of PrP^Sc^ structural heterogeneity within prion strains, suggesting the emergence and coevolution of structurally distinct PrP^Sc^ assemblies during prion replication in controlled environment. Such PrP^Sc^ diversification processes remain poorly understood. Although central to prion host-adaptation, structural diversification of PrP^Sc^ assemblies is also a key issue for the formation of PrP conformers involved in neuronal injury. Here, we characterized the evolution of the PrP^Sc^ quaternary structure during prion replication *in vivo* and in *bona fide* cell-free amplification assays. Regardless of the strain studied, the early replication stage conduced to the preferential formation of small PrP^Sc^ oligomers, thus highlighting a quaternary structural convergence phenomenon. Their evolutionary kinetics revealed the existence of a PrP^C^-dependent secondary templating pathway in concert with a structural rearrangement. This secondary templating pathway provides, for the first time, a mechanistic explanation for prion structural diversification during replication, a key determinant for prion adaptation on further transmission, including to other host species. The uncovered processes are also key for a better understanding of the accumulation mechanisms of other misfolded assemblies believed to propagate by a prion-like process.

## Introduction

In terms of pathogenic mechanisms, the prion paradigm unifies a number of age-related, incurable neurodegenerative disorders that are caused by protein misfolding and aggregation [13, 21, 22, 9]. These disorders include human and animal forms of prion diseases, Alzheimer’s disease, Parkinson’s disease and Huntington’s disease. In principle, host-encoded monomeric proteins or peptides are converted into misfolded and aggregated assemblies, which serve as seeds or templates for further autocatalytic conversion. In prion diseases, the ubiquitously expressed, host-encoded prion protein PrP^C^ is converted into a misfolded, β-sheet-rich conformer termed PrP^Sc^ [39]. In susceptible host species and in laboratory rodent models, PrP^Sc^ assemblies form stable, structurally distinct PrP^Sc^ conformers [4, 8, 12, 52], known as prion strains, and encode unique stereotypical biological phenotypes [45, 47, 48, 5]. The strain-specific structural differences can be observed at the secondary structural level in terms of local structural variation but also at the quaternary level with strain-specific size distributions [44, 49, 47]. A large body of evidence supports the view for further structural diversity within specific prion populations and strains: i) some studies highlight the selection of prion substrains during the transmission of natural isolates [11, 27, 2] or experimental prion strains [29] with a species or transmission barrier, ii) size- or density-fractionation studies support the existence of a heterogeneous spectrum of PrP^Sc^ assemblies with distinct tertiary/quaternary structures [24, 44, 49, 23, 50, 6, 7, 41] and biological activity (templating activity and infectivity) [24, 44, 49], and iii) kinetic studies of prion pathogenesis suggest that the formation of neurotoxic PrP^Sc^ species [46] occurs at the late stage of prion infection but that replicative PrP^Sc^ assemblies are formed at earlier stages [42, 43]. The prion replication process thus intrinsically allows the structural diversification of PrP^Sc^ assemblies.

While the kinetic aspects of prion replication ‘as a whole’ have been comprehensively described by measuring infectivity or PrP^Sc^ levels in the brain (see references [25, 34] as examples), the processes by which PrP^Sc^ structural diversification and the formation of different subpopulations occur within a given strain remain undescribed and are not mechanistically supported in the actual framework of the prion paradigm. The autocatalytic conversion model proposed by Griffith in 1967 [17], the nucleated-polymerization model described by Lansbury and Caughey in 1995 [26] and other derived models (e.g., [30]) merely assume the existence of structurally homogenous assemblies that have absolutely identical propensity to replicate throughout disease progression. These mechanisms intrinsically reduce PrP^Sc^ heterogeneity due to the best replicator selection process (*35, 36*). A recent high-resolution structural analysis of the N-terminal domain of the yeast prion SuP35 suggests that conformational fluctuations in natively disordered monomeric Sup35 are responsible for the stochastic, structural diversification of Sup35 aggregates [36]. This idea can be extrapolated to mammalian prion PrP to explain intrastrain structural diversification and strain mutation [12]. However, based on the best replicator selection concept [33, 37, 35], the aforementioned idea does not explain the coevolution of at least two structurally distinct PrP^Sc^ subassemblies within the same environment [11, 28].

To examine the molecular mechanisms of PrP^Sc^ replication and structural diversification in depth, we explored, with sedimentation velocity (SV)-based methods, the early stage of prion conversion *in vivo* and in a cell-free system by protein misfolding cyclic amplification (PMCA). PMCA mimics *in vivo* prion replication with accelerated kinetics [40]. By using several prion strains as templates, we demonstrated that the early stage of prion replication invariably generates two subsets of assemblies, termed A_i_ and B_i_, which differ in proportion, size and structure according to their specific infectivity. The analysis of their kinetics of formation during mb-PMCA combined with kinetic data assimilation revealed the existence of two sequential processes during prion replication. The first process corresponds to a quaternary structural convergence, as it tends to reduce the parental quaternary structure polydispersity to generate mostly small-sized assemblies, namely A_i_. The second process transforms the A_i_ into structurally different assemblies, namely, B_i_, according to a secondary auto-catalytic pathway requiring PrP^C^ and where B_i_ facilitates its own formation. Our findings provide, for the first time, mechanistic insights allowing the generation of structurally distinct assemblies during the prion replication process.

## Results

### Small PrP^Sc^ oligomers are formed at the early stage of prion replication

The early phases of prion replication are commonly thought to consist of an elongation/growing process [16], with the PrP^Sc^ template serving as a base. We studied the size distribution of proteinase K (PK)-resistant PrP^Sc^ (PrP^res^) assemblies at the early step of prion replication in the brain by SV in an iodixanol gradient using a previously published methodology [24, 49, 20]. The PrP^res^ size distribution at the disease end stage served as the control. Three different host PrP/strain combinations were studied: the 127S cloned scrapie prion strain in ovine PrP tg338 transgenic mice [25], the 139A cloned mouse prion strain in mouse PrP tga20 mice [15] and the vCJD cloned human prion strain in human PrP tg650 mice [3, 19]. As shown in Figure 1A-C, small oligomers sedimenting between fractions 1 and 4 were preferentially detected at the early stage of pathogenesis, regardless of the strain considered. A second population of oligomers with a larger size distribution and peaking in fractions 8-10 and 18 was observed for 127S. At the disease end stage and for the 3 strains, the small assemblies mostly disappeared at the expense of larger assemblies.

**Figure 1.**
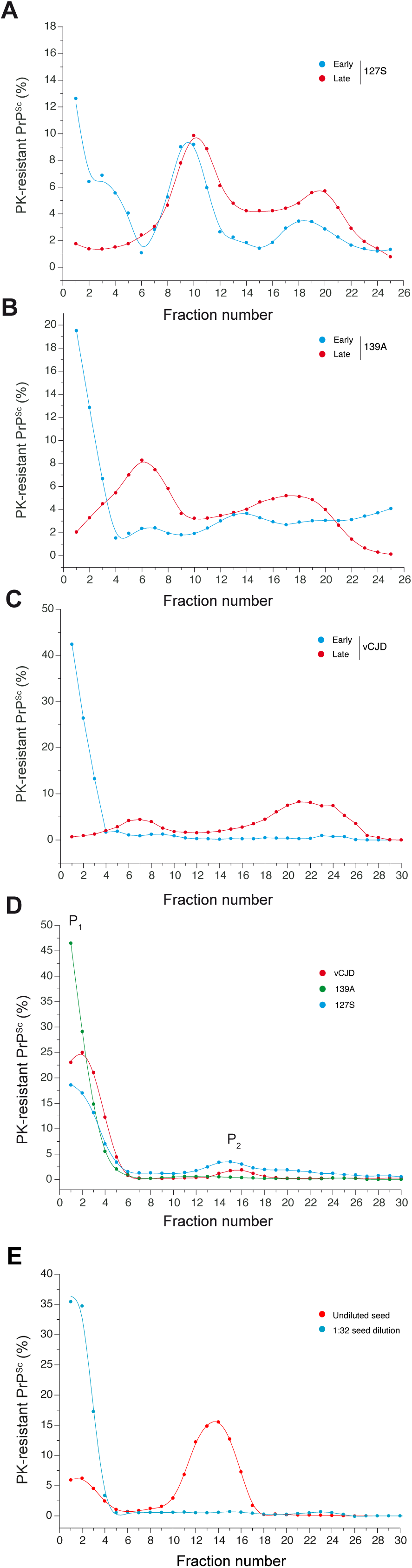
Size distribution of PrP^Sc^ assemblies from different prion strains at the early and late stages of pathogenesis *in vivo* and after the PMCA reaction. The size distribution of proteinase K (PK)-resistant PrP^Sc^ assemblies present in the brain *in vivo* (**A-C**) and in PMCA products (**D-E**) was examined by sedimentation velocity (SV). (**A-C**) For the *in vivo* sedimentograms, brains from ovine (tg338), murine (tga20) and human (tg650) transgenic mice inoculated with 127S scrapie prions (**A**), 139A mouse prions (**B**) and vCJD human prions (**C**) were collected (in triplicate) at the early stage (15 days postinfection (127S), 11 days postinfection (139A) and 120 days postinfection (vCJD), blue curves) and at the end stage of the disease (60 days postinfection (127S), 55 days postinfection (139A), 495 days postinfection (vCJD), red curves). The brains were solubilized and SV-fractionated. The collected fractions (numbered from top to bottom) were analyzed for PK-resistant PrP^Sc^ content by immunoblotting. (**D-E**) For the sedimentograms from the PMCA products with PrP^C^ substrate (**D**), the same strains were subjected to a single round of mb-PMCA by using 10^−5^ (139A) or 10^−6^ (vCJD, 127S) diluted brain homogenates as seed for the reaction. Thirty minutes after the last sonication, the amplified products were solubilized and SV-fractionated. The mean levels of PK-resistant PrP^Sc^ per fraction were obtained from the immunoblot analysis of *n=4* independent fractionations of PMCA reactions. The peaks containing PrP^Sc^ assemblies sedimentating in the top and middle fractions were termed P_1_ and P_2_, respectively. For the sedimentograms from the PMCA products without PrP^C^ substrate (**E**), undiluted 127S-infected tg338 brain (20% w/v, red curve) or a 1:32 dilution in PMCA buffer (blue curve) was used as seed, mixed with brain homogenate from PrP^0/0^ mice as substrate and subjected to a single round of mb-PMCA before SV fractionation (mean levels from *n=3* independent fractionations).

To determine whether the formation of small assemblies in the brain at the early stage of replication can be reproduced by an *in vitro bona fide* amplification method, we used a high-throughput variant of PMCA (i.e., mb-PMCA), which generates as much infectivity as in the brain at terminal stage of the disease in one unique round of 48 h, with high reproducibility in terms of limiting dilution and the amplification yield [31, 32]. When the size distribution of the amplified products was analyzed by SV after one mb-PMCA round, two discrete distributions were observed for the three strains (Figure 1D). The post-PMCA sedimentograms revealed the existence of a major set of small PrP^res^ assemblies sedimenting between fractions 1 and 3 (peak P_1_) and a minor set of larger assemblies with a well-defined Gaussian distribution centered on fraction 15 (peak P_2_). The relative proportions of P_1_ and P_2_ varied among the three strains; P_2_ was barely detected in the 139A amplicons. These data indicate that during mb-PMCA amplification, two populations of PrP^Sc^ assemblies are generated that differ according to their quaternary structures, with a predominance of small assemblies.

The bimodal (i.e., generation of two peaks) and discrete behavior of the size distribution as well as the formation of predominantly small assemblies in P_1_ suggest that the mb-PMCA condition(s) can be a consequence of shearing forces during the sonication step [1, 38, 51] rather than an intrinsic consequence of the replication process. To discriminate between these two possibilities, undiluted 127S seeds (i.e., 20% brain homogenate) were incubated and sonicated in identical mb-PMCA conditions but without the PrP^C^ substrate (i.e., in PrP^0/0^ brain lysate). As shown in Figure 1E, the size distribution analysis of these sonicated 127S seeds in the PrP^0/0^ substrate revealed mostly the presence of larger-sized assemblies, as observed upon solubilization at 37°C [24], thus ruling out the mb-PMCA conditions being at the origin of the formation of small-size assemblies.

Altogether, these observations suggest that *in vivo*, the early phase of replication for the 127S, 139A and vCJD prion strains generates mainly small-sized assemblies. Similar to *in vivo* replication, the mb-PMCA amplification condition generates two sets of PrP assemblies that differ in their quaternary structures. The formation of these two groups of assemblies is common to the three strains used here.

### P_1_ and P_2_ contain two structurally distinct PrP^res^ assemblies

We next asked whether the formation of P_2_ resulted from a simple condensation of assemblies present in the P_1_ peak (Oswald ripening process [53]) or from an alternative templating pathway. To address this question, we first examined the influence of the amplification rate on the formation of these two species by varying the concentration of the seed used as the template for the mb-PMCA reaction. We compared the SV-sedimentograms of the mb-PMCA products seeded with 10^−3^ to 10^−10^ diluted 127S brain homogenate. As shown in Figure 2A, as a function of the seed concentration, the relative amounts of assemblies in P_1_ decreased as the amounts of those from P_2_ increased. The variation in the P_1_ and P_2_ peak surface area as a function of the logarithm of the dilution factor revealed a quasi-linear decrease in P_1_ when the P_2_ peak surface followed a sigmoidal increase (Figure 2B). The sigmoidal increase in P_2_ to the detriment of the quasi-linear decrease in P_1_ surface indicates that i) the formation of PrP^res^ assemblies present in P_2_ follows a seed concentration-dependent cooperative process and that ii) the formation of the P_2_ peak does not result from the simple condensation of assemblies present in P_1_ as the variations in P_1_ and P_2_ are uncorrelated (Figure 2B). This observation strongly suggests that assemblies forming the P_1_ and P_2_ peaks result from distinct polymerization pathways and should therefore be structurally distinct.

**Figure 2.**
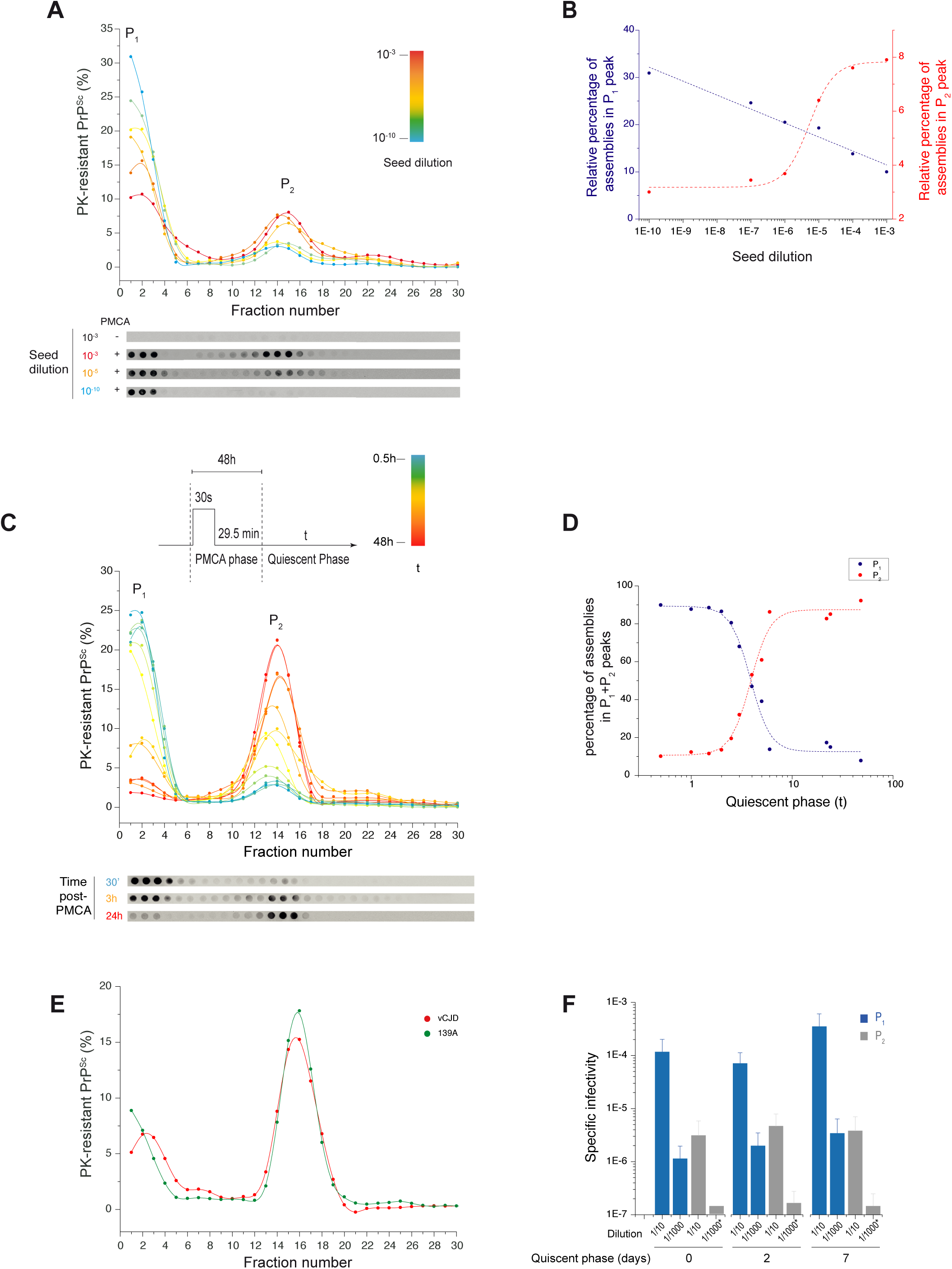
Seed concentration- and time-dependent dynamic evolution of the PMCA-generated PrP^Sc^ assemblies. (**A-B**) SV profile of mb-PMCA products seeded with ten-fold dilutions from 127S-infected brain homogenates, as indicated. Thirty minutes after the last sonication, the amplified products were solubilized and SV-fractionated. The mean relative levels of PK-resistant PrP^Sc^ per fraction (**A**) were obtained from the immunoblot analysis of *n=4* independent fractionations of PMCA reactions (representative dot-blot shown). Variation in the P_1_ and P_2_ peak surface areas as a function of the logarithm of the seed dilution factor (**B**). **(C)** PK-resistant PrP^Sc^ sedimentograms from the PMCA products generated with 127S prions (10^−5^ dilution) and further incubated at 37°C during the indicated quiescent phase (t), i.e., without sonication. At each time point, the collected products were frozen. All collected samples were then thawed, fractionated in parallel by SV and analyzed by immunoblot (**C**, *n=3* independent experiments, representative dot-blot shown). **(D)** PK-resistant PrP^Sc^ isopycnic sedimentograms from PMCA products generated with 127S prions (10^−5^ dilution) and immediately fractionated at the end of the PMCA reaction (blue line and symbol) or after a 24h-quiecent incubation at 37°C (red line and symbol). At each time point, the collected samples were frozen. All collected samples were then thawed, fractionated in parallel by sedimentation at the equilibrium [24] and analyzed by immunoblot (the mean levels of PK-resistant PrP^Sc^ per fraction were obtained from the immunoblot analysis of *n=3* independent fractionations of PMCA reactions). As control, the density profile of PK-resistant PrP^Sc^ assemblies from the brain of terminally sick tg338 mice (solubilization at 37°C to mimic the PMCA conditions) is shown (gray line and symbol). **(E)** Evolution of the percentage of P_1_ and P_2_ peaks as a function of the quiescent phase post-PMCA reaction (**C**). **(F)** PK-resistant PrP^Sc^ sedimentograms from the PMCA products generated with 139A and vCJD prion seeds (10^−5^ dilution) and further incubated for a quiescent period of 24 h at 37°C. **(G)** Specific infectivity of the P_1_ and P_2_ peaks post-PMCA reaction and after quiescent incubation. Fractions corresponding to P_1_ (fractions 1-3) and P_2_ (fractions 14-16 (days 0 and 2) or 16-18 (day 7)) were pooled and inoculated into groups of reporter tg338 mice at two different dilutions (1:10 and 1:1000) for better accuracy. The specific infectivity of the assemblies was calculated from the mean survival time of the mice using a 127S dose-response curve. *: incomplete attack rate.

To further explore the entanglement between the assemblies forming P_1_ and P_2_, we fixed the mb-PMCA regime to favor the formation of the P_1_ peak by fixing high dilutions of the inoculum seed, followed by quiescent incubations at 37°C for variable periods. As shown with the 127S prions, the SV analysis at defined incubation time points post-PMCA reaction revealed a decrease in the population of P_1_ in favor of P_2_ (Figure 2C). At 4 h postincubation, there were equal proportions of assemblies forming P_1_ and P_2_. At 24 h, most of the PrP^res^ assemblies were located in the P_2_ peak. Comparing the distribution in isopycnic gradients [24] of the PrP^res^ populations at 0h and 24h of quiescent incubation at 37°C revealed a quasi-similar density for PrP^res^ assemblies composing the P_1_ and the P_2_ peaks (Figure 2D). This observation leads us to conclude that i) the low sedimentation velocity of the assemblies forming P_1_ does not result from an interaction with lipids or other low-density molecules and ii) the sedimentation velocity increase of P_2_ compared to P_1_ results strictly from a quaternary structure rearrangement through size increase rather than change in compactness reducing the hydrodynamic radius.

As shown in Figure 2E, the formation of assemblies sedimenting in P_2_ exhibited bimodal behavior (i.e., absence of assemblies of intermediate size) without any significant shift in the P_2_ peak position, suggesting that the formation of these assemblies resulted from the association with a specific number of assemblies present in P_1_. Furthermore, the time-dependent surface variation in P_1_ and P_2_ showed a sigmoidal shape, indicating that the assemblies present in P_2_ enhance their own formation according to an autocatalytic process (Figure 2E). Similarly, the 139A and vCJD prions showed a bimodal evolution of P_1_ to P_2_ during a 24-h quiescent phase (Figure 2F), arguing in favor of a generic process of transformation.

To determine whether the quaternary structure rearrangement accompanying the transformation of P_1_ to P_2_ was in concert with a deeper structural rearrangement in the PrP^Sc^ assemblies, we determined the specific infectivity of the P_1_ and P_2_ assemblies. A 127S-PMCA product was fractionated at the end of the reaction or after 48 h of quiescent incubation. Pools of fractions corresponding to the P_1_ and P_2_ peaks were inoculated into reporter tg338 mice. The specific infectivity (infectivity per PrP molecule), which is mostly associated to PrP^res^ assemblies [24, 49], was calculated from the mean survival time using 127S dose-response curves [49]. As shown in Figure 2G, the specific infectivity of the P_1_ peak assemblies was 50-100-fold higher than that of the P_2_ peak assemblies. This value did not change over a longer period of quiescent incubation (7 days, Figure 2G).

To determine whether the P_2_ peak assemblies could further evolve, we extended the quiescent phase up to 30 days. For the 127S, 139A and vCJD prion strains, the sedimentogram curves at 7 and 30 days showed a translational shift in the P_2_ peak to higher fractions, indicative of an isokinetic increase in their mean average sizes (Figure 3, left curves). The difference in the specific infectivity values of the P_1_ and P_2_ peak assemblies did not change over a longer period of quiescent incubation (7 days, Figure 2G, 127S strain). This size translation thus contrasts with the bimodal phase and highlights a change in the kinetic regime.

**Figure 3.**
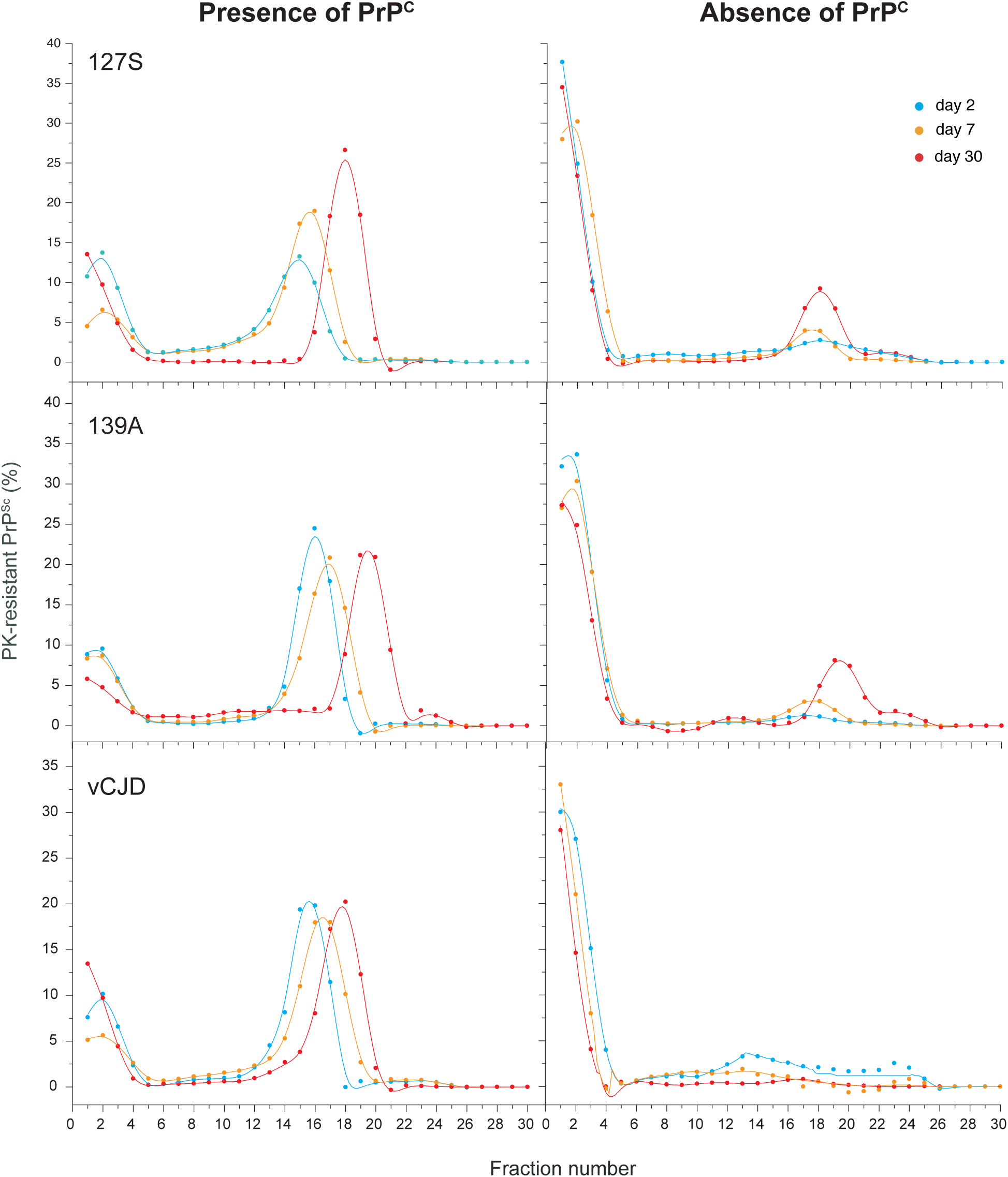
PrP-dependent generation of B_i_ assemblies from A_i_ assemblies. PMCA products from 127S, 139A and vCJD prions (10^5^, 10^4^ and 10^4^ diluted seeds, respectively) were treated with or without PK to eliminate PrP^C^ before quiescent incubation at 37°C for 24 h, 7 days or 30 days, as indicated. At each time point, the collected products were frozen. All collected samples were then thawed, SV-fractionated in parallel and analyzed by immunoblotting (*n=3* independent experiments).

Collectively, these observations indicate that the P_1_ and P_2_ peaks contain structurally distinct sets of PrP^res^ assemblies named A_i_ and B_i_. The index i indicates their sizes, with A_i_ and B_i_ being the major constituents of the P_1_ and P_2_ peaks, respectively. The formation of the B_i_ assemblies is cooperative and results from a complex kinetic pathway. Upon longer quiescent incubation, a change in the kinetic regime occurs.

### The formation of B_i_ from A_i_ assemblies requires the presence of PrP^C^ during the quiescent phase

Our previous studies revealed that only ∼30% of the PrP^C^ substrate was converted into PrP^Sc^ after a complete round of mb-PMCA [31, 32]. To determine whether the remaining 70% still participated in the transformation of A_i_ to B_i_ assemblies during the quiescent phase, PMCA products from the 139A, 127S and vCJD prions were treated with PK to eliminate PrP^C^ before quiescent incubation at 37°C. As shown in Figure 3 (right curves), the amount of B_i_ assemblies generated during the 48-h quiescent incubation was drastically decreased for the three prion strains in the absence of the PrP^C^ substrate. Further quiescent incubation for 7 and 30 days in the absence of PrP^C^ allowed the formation of low amounts of B_i_ assemblies for the 127S and 139A prion strains. The fact that the transformation of A_i_ to B_i_ assemblies is strongly facilitated by the presence of PrP^C^ suggests that B_i_ assemblies result from the integration/conversion of PrP^C^ into A_i_ assemblies. The appearance of a low amount of B_i_ after a long incubation period without PrP^C^ may result from the leakage of monomers from a conformer cosedimenting with A_i_.

### Kinetical scheme describing the transformation of A_i_ to B_i_ assemblies

To establish a kinetic mechanism and provide a molecular interpretation of the assemblies’ dynamics during the quiescent phase, a number of elementary steps were identified based on experimental observations and were used as unavoidable constraints [14]. The first constraint was the existence of two structurally distinct PrP^Sc^ subassemblies, namely, A_i_ and B_i,_ with distinct dynamics. Indeed, structurally equivalent assemblies would fail to present a bimodal size distribution, cooperative seed concentration and kinetic evolution or distinct specific infectivity. The second constraint was the existence of a detailed balanced between the PrP^Sc^ assemblies and their elementary subunit (suPrP), as previously shown [20] making the size distribution of the PrP^Sc^ assemblies highly dynamic and dependent on the assembly concentration, as shown in Figure 1E. Indeed, SV analysis of the PrP^0/0^ brains lysates seeded with 30-fold-diluted 127S-infected brains and submitted to PMCA revealed a quaternary structure rearrangement with a shift in lower molecular weight assemblies according to the detailed balance:

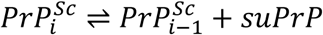

where 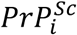 and 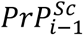 are the sizes *i* and *i-1* of suPrP^Sc^, respectively.

Because the existence of suPrP is a generic property of prion strains [20], the 3^rd^ constraint leads us to assume that the A_i_ and B_i_ assemblies are in detailed balance with their respective suPrPs (denoted *suPrP*^*A*^ and *suPrP*^*B*^) but with distinct equilibrium constants 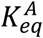 and 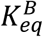. Thus, at any moment of the process of assembly transformation of A_i_ to B_i_, the following equilibrium should be respected:

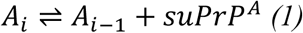

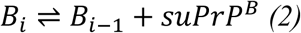

The equilibrium constant 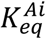 and 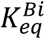 governs the respective size distribution of the A_i_ and B_i_ assemblies and, thus, the bimodal aspect of the curve. According to our previous SV calibrations with PrP oligomers and globular mass markers [49], the size distribution of the A_i_ and B_i_ subassemblies were fixed: *i*_*A*_<5 and *i*_*B*_ centered around 20 PrP-mers. Due to the limited resolution of SV fractionation for small assemblies, we assumed that *A*_*i*_ and suPrP^B^ cosedimented. The fourth constraint relies on the fact that the transformation of A to B requires PrP^C^ and that the kinetic is cooperative, as shown in Figures 1E and 2. This cooperativity implies that B subassemblies facilitate their own formation according to an autocatalytic process. This can be resumed by the following minimalistic autocatalytic process:

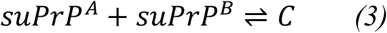

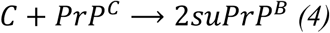

where C is an active complex reacting with PrP^C^ that generates B assemblies. Considering that suPrP^B^ can condense into B_2_ [20] and according to detailed balance (2), one can establish the reaction model describing the formation of B_i_ assemblies from the neo-formed *suPrP*^*B*^:

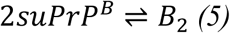

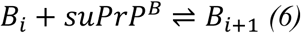

Altogether, these six elementary steps constitute the reaction mechanism that describes the transformation of A_i_ into B_i_ subassembly species.

To validate the designed mechanism, we translated these elementary reactions into time-dependent differential equations (for more details, see Supplementary text) and performed kinetic simulations using the size distribution of the PrP^Sc^ assemblies immediately after cyclic amplification as the initial condition (blue curve in Figure 2A). According to the model, the simulated size distribution variation as a function of time showed bimodal behavior, as was experimentally observed (Figure 4A). Furthermore, the theoretical size distribution centroid presented similar sigmoidal patterns to those of the experimental data (Figure 4B), arguing in favor of an autocatalytic kinetic model describing the overall quaternary structure evolution of PrP^Sc^ assemblies during the quiescent phase. The analysis of the model (for more details, see Supplementary text) revealed that the autocatalytic formation of B_i_ species occurs at the expense of A_i_ species and with PrP^C^ consumption (Figure 4 C and D). According to this model, when PrP^C^ is in large excess, A_i_ constitutes the limiting compound for the formation of B_i_ assemblies. Therefore, during the quiescent phase, the PrP^C^ to PrP^Sc^ conversion rate is directly proportional to the amount of A_i_ assemblies (Figure 4D).

**Figure 4.**
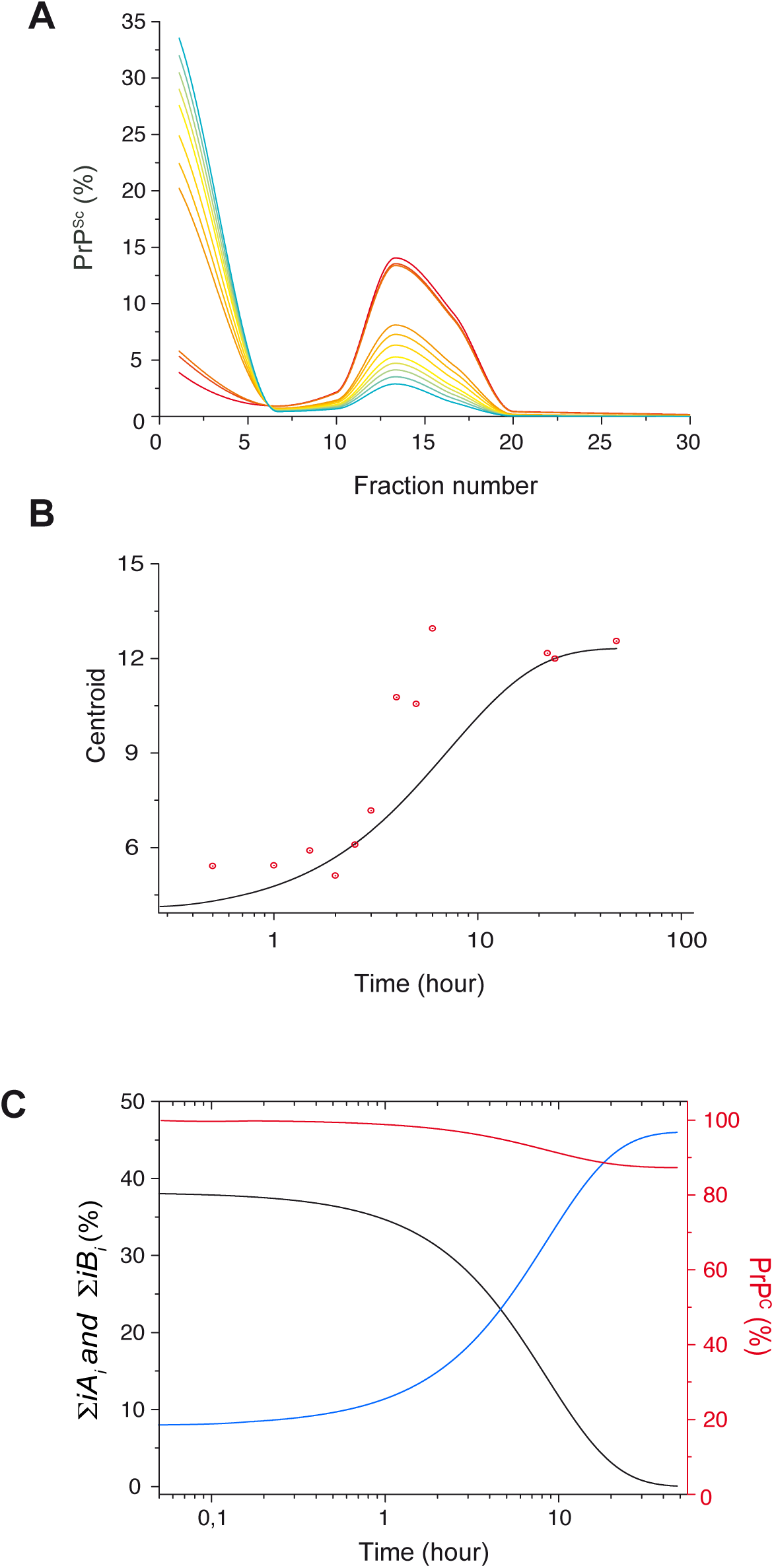
Mathematical modeling of the time-dependent dynamic evolution of the PMCA-generated PrP^Sc^ assemblies. **(A)** The size distribution evolution of a structurally distinct set of assemblies A_i_ and B_i_ dimensioned on gradient fraction numbers was simulated based on the kinetic scheme described in the results section (equations 1 to 6) and the Supplementary text. **(B)** The time dependency evolution of the simulated centroid (black line) and centroid calculated from experimental sedimentograms of Figure 2D (red circle) show a similar shape, supporting the cooperativity hypothesis of the transformation of A_i_ into B_i_. (**C**) The simulation of time dependency evolution of the total amount of A_i_ assemblies (Σ *iA*_*i*_ in black), B_i_ assemblies (Σ *iB*_*i*_ in blue) and the monomer (in red) revealed that A_i_ assemblies constitute the limiting species for the conversion of PrP^C^ during the quiescent phase. In the present simulation framework (for more details, see Supplementary text), only 14% of PrP^C^ is consumed.

## Discussion

The mechanisms of prion replication and the dynamics responsible for prion structural diversification in the infected host remain unclear and rarely addressed. In the actual framework of the prion paradigm, the templating process is admitted to occur at the prion assembly interface, leading to an increased size of the complex formed by the template:substrate, out of the fragmentation/depolymerization context. The atypical size distribution observed here at the early stage of the replication process for three distinct prion strains, where accumulation of small-size assemblies dominates, contrasts with this canonical templating model and requires an additional process that considers the dynamics of replication. Furthermore, the existence of a multistep conversion process provides an unexpected approach to reconciling the best replicator [35] selection paradigm and diversification process, which is inherent to prion adaptation and evolution.

As shown *in vivo* for the vCJD, 127S and 139A prion strains, the early stage of the replication process in the brain is dominated by the accumulation of small assemblies, whereas higher-size subsets are mostly detected at the terminal stage of pathogenesis. Such quaternary structural diversity, - and beyond the existence of structurally distinct types of assemblies, as defined by their specific infectivity ([24, 49] and Supplemental Figure 1), can be exclusively explained by the existence of a balance between at least two kinetic modes taking place at different stages of the pathogenesis. Both can be governed by evolution or a fluctuation in the replication microenvironment due to the physio-pathological state of the infected animal and/or to the sequential involvement of specific prion-replicating cell types. However, another possibility can lie in the intrinsic and deterministic properties of the PrP replication process to generate structurally distinct types of assemblies. Discriminating between these two nonmutually exclusive hypotheses is technically difficult *in vivo*. The mb-PMCA as a *bona fide* amplification method in a more simplified and kinetically controlled context constitutes a relevant method for investigating the intrinsic propensity of the replication process to generate structurally distinct assemblies. Interestingly, and against common belief, the size distribution of the PrP^Sc^ assemblies used as seeds was relatively insensitive to mb-PMCA sonication cycles when a simple dilution displaced the assemblies towards a smaller size (Figure 1E), as previously reported [20]. These two observations exclude the contribution of the fragmentation process during the mb-PMCA sonication cycles to the size distribution pattern of PrP^Sc^ assemblies and emphasize the existence of a constitutional dynamic between the PrP^Sc^ subpopulation [20], which should be considered during the replication process. We showed that two sets of PrP^Sc^ assemblies, A_i_ and B_i,_ were generated during the mb-PMCA reaction. The A_i_ and B_i_ assemblies constitute two structurally distinct PrP^Sc^ subpopulations, as supported by their distinct specific infectivity; the bimodal size distribution instead of a continuum; the effect of initial seed concentration on the respective proportions of A_i_ and B_i_; and the role of PrP^C^ in the transformation of A_i_ to B_i_ that indicates that B_i_ assemblies do not result from simple condensation of A_i_ assemblies. Therefore, the prion replication process *per se* intrinsically generates structurally diverse PrP^Sc^ subassemblies.

According to our SV experiments, small-sized PrP^Sc^ assemblies were mainly formed at the early stage of prion replication in the brain and during the mb-PMCA reaction. This was observed with three distinct prion strains (127S, 139A, vCJD) on 3 different PrP genetic backgrounds. Considering that the PrP^Sc^ assemblies that constitute each strain are structurally distinct, one can ask how distinct PrP^Sc^ assemblies can all generate A_i_ assemblies that harbor strain structural information while showing the same quaternary structure (at the SV resolution). The first explanation can be the existence of a common narrow subpopulation of PrP^Sc^ (with respect to their quaternary structure) within the three strains that serves as the best replicator and participates in the formation of A_i_ assemblies. However, the PrP^Sc^ quaternary structure subset that exhibits the highest specific infectivity *in vivo* (i.e., the best replicator) can be associated with either small-size assemblies (i.e., 127S and 139A [24, 49] and Supplemental Figure 1A, respectively) or high-molecular-weight assemblies (i.e., vCJD, Supplemental Figure 1B) and is therefore strain-dependent. The existence of a structurally common PrP^Sc^ subpopulation is thus unlikely to be at the origin of the generic formation of a small-size subset in the brain or A_i_ assemblies in the mb-PMCA condition. Intrinsically, the early steps of the replication process favor the emergence of mainly one subspecies A_i_ with a highly narrowed size distribution, arguing in favor of a quaternary structural convergence phenomenon during these steps. This structural convergence concerns the PrP domain that governs polymerization (the size of assemblies). However, as the A assemblies harbor the strain structural determinant, one can conclude that A_i_ assemblies present a certain degree of structural variability, allowing the encoding of strain structural information (Supplemental Figure 1).

All along the quiescent phase and for the three prion strains studied, the A_i_ assemblies constitute the precursor species in the formation of B_i_ assemblies. Furthermore, the presence of PrP^C^ is required for the evolution of A_i_ into B_i_ assemblies, and according to the kinetic model describing the autocatalytic formation of B_i_ during the quiescent phase, A_i_ is the limiting species for conversion when large amounts of PrP^C^ are present (Figure 4C and D and Supplementary text). The bimodal quaternary structure evolution during the quiescent phase is in concert with a specific infectivity decrease that is indicative of a structural rearrangement of species present in P_1_ and P_2_ and thus during the transformation of A_i_ to B_i_. Even if the first event conducing to the formation of B_i_ assemblies remains undetermined, we can assume that A_i_ can have the intrinsic propensity to spontaneously evolve into B_i_ assemblies in the presence of PrP^C^ (Figure 5). The cooperative disappearance of P_1_ in favor of P_2_ strongly suggests an autocatalytic process for the transformation of A_i_ to B_i_ (reactions 3 and 4). This last phenomenon shows the existence of a secondary autocatalytic process, undescribed until now, in the canonical prion replication process [26]. One can be reasonably envisage that A_i_ can have the intrinsic propensity to generate B_i_ assemblies in the presence of PrP^C^ assemblies with a very low efficiency. This parallel pathway to the autocatalytic process can then explain how the first set of B_i_ assemblies is generated (Figure 5). The existence of a secondary autocatalytic process can be crucially important for maintaining PrP^Sc^ structural diversity throughout the evolution of the pathology. In the absence of this secondary autocatalytic process, e.g., in the absence of PrP^C^, the system selects the best replicator and the most thermodynamically stable assemblies. In the presence of PrP^C^, the system escapes this rule, allowing the specific accumulation of the autocatalytic product (here, the B_i_ assemblies) rather than the assemblies that are the most thermodynamically stable or have highly specific infectivity. This phenomenon can explain why, for certain prion strains, the most infectious assemblies represent a minor population, while those with the lowest specific infectivity mostly accumulate [24, 49].

**Figure 5.**
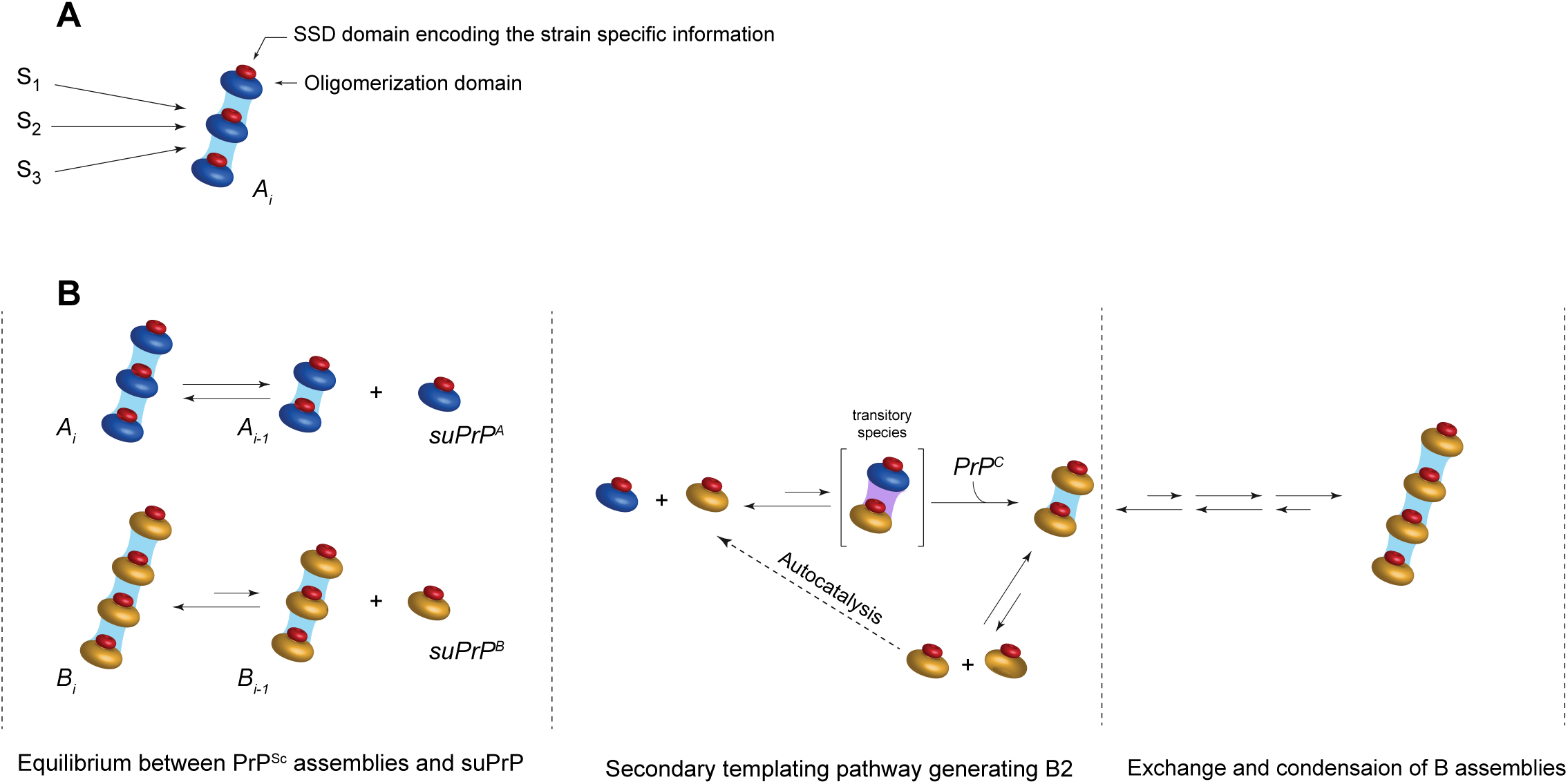
Quaternary structural convergence and secondary autocatalytic pathway at the root of the formation of B_i_ assemblies. **(A)** Different prion strains (S_1_, S_2_ and S_3_) give rise to the formation of common oligomeric assemblies, termed A_i_, with a narrowed size distribution during mb-PMCA reactions. This common quaternary structural convergence at the early stage of the replication process suggests the existence of a common conversion pathway and a common oligomerization domain that is independent of the strain structural determinant (SSD, represented in red). **(B)** According to the dilution experiments (see Figure 1E), an equilibrium exists between PrP^Sc^ assemblies and the suPrP from each subpopulation [20]. Based on the constraints imposed by the experimental observations, the best model to account for the cooperative and PrP^C^ dependency transformation of A_i_ into B_i_ assemblies implicates a secondary templating pathway where the transformation of suPrP^A^ to suPrP^B^ is assisted by suPrP^B^, rendering the process autocatalytic.

## Conclusion

The early step of prion replication for at least three distinct prion strains leads to the formation of small assemblies. The mb-PMCA approach clearly demonstrates the intrinsic properties of the *bona fide* replication process to generate at least two structurally distinct PrP^Sc^ subassemblies. The deterministic aspect of the replication process to generate a structurally diverse set of assemblies contrasts with the widespread idea that considers the prion diversification process within a given strain (often referred to as the creation of prion quasi-species) as a stochastic event and as a process that is cogoverned by environmental fluctuations [52]. The secondary autocatalytic pathway leading to the formation of B_i_ subassemblies can participate in prion adaptation during transmission events with species barriers. Considering that the transmitted inoculum initially contains A_i_ and B_i_ assemblies, the autocatalytic conversion process of B_i_ can kinetically drive the adjustment and integration of the new-host PrP^C^ to generate host-adapted B_i_ assemblies. This hypothesis is supported by our recent observations in which complementation between A_i_ and B_i_ subassemblies is required to cross existing species barriers (submitted article).

## Methods

### Ethics

Animal experiments were conducted in strict accordance with ECC and EU directives 86/009 and 2010/63 and were approved by the local ethics committee of the author’s institution (Comethea; permit numbers 12/034 and 15/045).

### Transgenic mouse lines and prion strains

The ovine (tg338 line; Val136-Arg154-Gln171 VRQ allele), human (tg650 line; Met129 allele) and mouse (tga20) PrP transgenic lines have been described previously [3, 15, 25]. The mouse lines were homozygous and overexpressed approximately 8-, 6-, and 10-fold amounts of heterologous PrP^C^ on a mouse PrP-null background. PrP^0/0^ mice were the so-called Zürich-I mice [10]. Cloned 127S scrapie, human vCJD and mouse 139A prion strains were serially passaged in tg338, tg650 and tga20 mice, respectively [31, 32]. These strains were used as pools of mouse-infected brains and prepared as 20% wt/vol homogenates in 5% glucose by use of a tissue homogenizer (Precellys 24 Ribolyzer; Ozyme, France).

### Time course analysis of prion accumulation

Eight-week-old female tg338, tg650 and tg20 mice were inoculated intracerebrally in the right cerebral hemisphere with 127S, vCJD or 139A prions (20 μl of a 10% brain homogenate dose). Infected animals were euthanized by cervical column disruption in triplicate at regular time points and at the terminal stage of disease. Brains were removed and kept for PrP^Sc^ size fractionation.

### Miniaturized bead-PMCA assay

The miniaturized bead-PMCA assay [11, 20, 32] was used to amplify prions. Briefly, serial ten-fold dilutions of 127S, vCJD and 139A prions (mouse brain homogenates diluted in PMCA buffer) were mixed with brain lysates (10% wt/vol) from healthy tg338, tg650 and tga20 mice as respective substrates and subjected to one round of 96 cycles of 30-s sonications (220-240 Watts) followed by 29.5 min of incubation at 37°C. With a >10^4^ dilution of the seeds, input PrP^Sc^ is not detected in the mb-PMCA products. PMCA was performed in a 96-well microplate format using a Q700 sonicator (QSonica, USA, Delta Labo, Colombelles, France). For quiescent incubation, the samples were left in the incubator at 37°C for the indicated period of time, without any sonication. To eliminate residual PrP^C^ present in the PMCA products before quiescent incubation, the samples were treated with PK (80 μg/ml final concentration). The treatment was stopped by adding 2 mM Pefabloc and 1x EDTA-free protease inhibitor cocktail. All final products were kept for PrP^Sc^ size fractionation, and aliquots were PK-digested (115 μg/ml final concentration, 0.6% SDS, 1 h, 37°C) prior to immunoblot analyses, as described below.

### Sedimentation velocity fractionation

SV experiments were performed as described previously [24, 49, 20]. Mouse brain homogenates or PMCA products were solubilized by adding an equal volume of solubilization buffer (50 mM HEPES pH 7.4, 300 mM NaCl, 10 mM EDTA, 2 mM DTT, 4% wt/vol dodecyl-β-D-maltoside (Sigma)) and incubated for 45 min on ice. Sarkosyl (N-lauryl sarcosine; Fluka) was added to a final concentration of 2% wt/vol, and the incubation continued for an additional 30 min on ice. A total of 150 μl of solubilized samples was loaded atop a 4.8-ml continuous 10– 25% iodixanol gradient (Optiprep, Axys-Shield), with a final concentration of 25 mM HEPES pH 7.4, 150 mM NaCl, 2 mM EDTA, 1 mM DTT, 0.5% Sarkosyl. The gradients were centrifuged at 285,000 g for 45 min in a swinging-bucket SW-55 rotor using an Optima LE-80K ultracentrifuge (Beckman Coulter). Gradients were then manually segregated into 30 equal fractions of 165 μl from the bottom using a peristaltic pump and analyzed by immunoblotting or bioassay for PrP^Sc^ or infectivity, respectively. To avoid any cross-contamination, each piece of equipment was thoroughly decontaminated with 5 M NaOH followed by several rinses in deionized water after each gradient collection [24].

### Isopycnic sedimentation

The entire procedure was performed as described previously [24]. Mouse brain homogenates or PMCA products were solubilized as described above. For mouse brain homogenates, solubilization was performed at 37°C to mimic PMCA conditions. A total of 220 μl of solubilized material was mixed to reach 40% iodixanol, 25 mM HEPES pH 7.4, 150 mM NaCl, 2 mM EDTA, 1 mM DTT, 0.5% Sarkosyl final concentration and loaded within a 4.8 ml of 10– 60% discontinuous iodixanol gradient with a final concentration of 25 mM HEPES pH 7.4, 150 mM NaCl, 2 mM EDTA, 1 mM DTT, 0.5% Sarkosyl. The gradients were centrifuged at 115 000 g for 17 hours in a swinging-bucket SW-55 rotor using an Optima LE-80K ultracentrifuge (Beckman Coulter). Gradients were then manually segregated into 30 equal fractions of 165 μl from the bottom using a peristaltic pump and analyzed for PrP^Sc^ content by immunoblotting.

### Analysis of PrP^Sc^ content by immunoblotting

Aliquots of the SV-fractionated PMCA samples were treated with PK (50 μg/ml final concentration, 1 h, 37°C) before mixing in Laemmli buffer and denaturation at 100°C for 5 min. The samples were run on 12% Bis-Tris Criterion gels (Bio-Rad, Marne la Vallée, France) and electrotransferred onto nitrocellulose membranes. In some instances, denatured samples were spotted onto nitrocellulose membranes using a dot-blot apparatus (Schleicher & Schuell BioScience (Whatman)). Nitrocellulose membranes were probed for PrP with 0.1 μg/ml biotinylated anti-PrP monoclonal antibody Sha31. Immunoreactivity was visualized by chemiluminescence (GE Healthcare). The protein levels were quantified with ImageLab software after acquisition of chemiluminescent signals with a Chemidoc digital imager (Bio-Rad, Marnes-la-Coquette, France). For all SDS-PAGE analyses, a fixed quantity of human recombinant PrP was employed for consistent calibration of the PrP signals in different gels.

### Bioassays

The pool of fractions of interest was extemporarily diluted ten-fold in 5% glucose and immediately inoculated via the intracerebral route into reporter tg338 mice (20 μl per pool of fraction, n = 5 mice per pool). Mice showing prion-specific neurological signs were euthanized at the end stage. To confirm prion disease, brains were removed and analyzed for PrP^Sc^ content using the Bio-Rad TsSeE detection kit [27] prior to immunoblotting, as described above. The survival time was defined as the number of days from inoculation to euthanasia. To estimate what the difference in mean survival times means in terms of infectivity, strain-specific curves correlating the relative infectious dose to survival times were used, as previously described [49].

### Kinetic simulation

The details of the kinetic simulation are reported in the Supplementary text. Briefly, two distinct sets of assemblies were considered (A_i_ and B_i_). Based on experimental observations, a set of constraints was retained to build biochemical reactions describing the evolution of the quaternary structure of PrP^res^ assemblies. The ordinary differential equations of the biochemical reactions 1 to 6 (in the manuscript) were established and coded in MATLAB for simulations.

**Supplemental Figure 1. PK-resistant PrP^Sc^ and the infectivity sedimentation profile of 139A and vCJD prion strains**

Brain homogenates from tga20 mice infected with 139A prions (**A**) and tg650 mice infected with vCJD prions (**B**) were solubilized and SV-fractionated. The collected fractions were analyzed for PK-resistant PrP^Sc^ content (black line) and for infectivity (red bars or line) with an incubation time bioassay in reporter tga20 and tg650 mice. The mean survival time values of these mice were reported as standard dose-response curves ([18] and unpublished) to determine relative infectious dose values. A relative infectious dose of 0 corresponds to animals inoculated with 2 mg of infectious brain tissue.

## Acknowledgments

We thank the staff of the Animal facility (INRA-UEAR, Jouy-en-Josas) for animal care. This work was supported by grants from the Fondation pour la Recherche Médicale (Equipe FRM DEQ20150331689), the European Research Council (ERC Starting Grant SKIPPERAD, number 306321), and the Ile de France region (DIM MALINF).

