## Supplemental material for "Quaternary structural convergence and structural diversification of prion assemblies at the early replication stage"

#### Supplementary material

- Supplementary figure 1
- Supplementary text

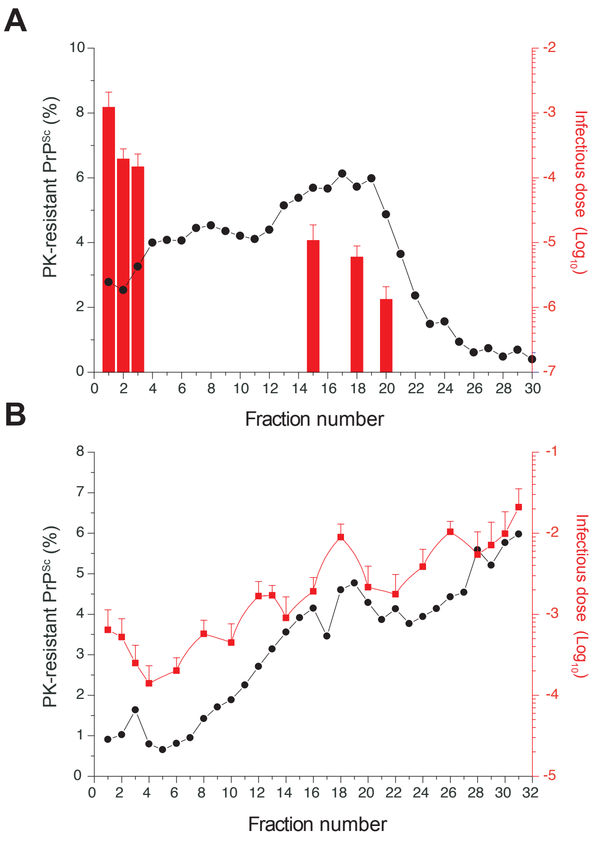


**Supplemental Figure 1. PK-resistant PrP$^{Sc}$ and the infectivity sedimentation profile of 139A and vCJD prion strains**

Brain homogenates from tga20 mice infected with 139A prions (**A**) and tg650 mice infected with vCJD prions (**B**) were solubilized and SV-fractionated. The collected fractions were analyzed for PK-resistant PrP$^{Sc}$ content (black line) and for infectivity (red bars or line) with an incubation time bioassay in reporter tga20 and tg650 mice. The mean survival time values of these mice were reported as standard dose-response curves ([18] and unpublished) to determine relative infectious dose values. A relative infectious dose of 0 corresponds to animals inoculated with 2 mg of infectious brain tissue.

### Supplementary Information: Mathematical modelling

November 21, 2018

#### 1 Reaction scheme

As explained in the Main Text, we consider two different kinds of oligomers: on the one hand, $A_i$, of size $2i$, are formed by the aggregation of $i$ SuPrP formed of two monomers, and denoted $A_1$. Due to the fact that $i_A < 5$, as $A_i$ assemblies are eluded in the first Sedimentation Velocity (S.V) fractions, we neglect here the oligomers $A_i$ with $i > 1$ for the sake of clarity and simplicity. On the second hand, oligomers $B_i$, of size $3i$, able to aggregate by $B_1-$ addition, where $B_1$ is another SuPrP formed of three monomers. However, $A_1$ may react with monomers to give rise to $B_1$. Let us also note that the size of $A_1$ and $B_1$, respectively formed of two and three monomers, is somewhat arbitrary: all we know is that this is their order of magnitude, in coherence with [1]. As explained in the Main Text, a convenient reaction scheme should also be such that without monomers, almost nothing happens (Fig. 3 in the Main Text).

1. $A_1$ and $B_1$ can form a complex $C$ in a reversible way:

$$A_1 + B_1 \underset{k_C^-}{\overset{k_C^+}{\rightleftharpoons}} C.$$

2. The complex $C$ can then react with the monomer $M$ to form two $B_1$:

$$M + C \xrightarrow{k_+} 2B_1.$$

3. The oligomers $B_i$ follow a classical polymerisation/depolymerisation chain reaction, by $B_1-$ addition:

$$B_1 + B_i \underset{k_{dep}^{i+1}}{\overset{k_{on}^i}{\rightleftharpoons}} B_{i+1}, \qquad 1 \le i \le n-1.$$

#### 1.1 Ordinary differential equations

The data obtained by S.V are interpreted as a dilatation of a size distribution density, normalized at 100%, so that if $u_i$ denotes the concentration of polymers formed of $i$ monomers, the data represent $\frac{iu_i}{\sum\limits_k ku_k}$.

We translate this reaction scheme into the following system of differential equations, denoting $a_1$, $b_1$, $c$ and $b_i$ respectively the concentrations of $A_1$, $B_1$, $C$ and $B_i$ :

$$\frac{dm}{dt} = -k_+ m(t)c(t), \qquad m(0) = m^0, \tag{1.1}$$

$$\frac{dc}{dt} = k_C^+ a_1(t)b_1(t) - k_C^- c(t) - k_+ m(t)c(t), \qquad c(0) = c^0, \tag{1.2}$$

$$\frac{da_1}{dt} = -k_C^+ a_1(t)b_1(t) + k_C^- c(t), \qquad a_1(0) = a_1^0, \tag{1.3}$$

$$\frac{db_1}{dt} = -k_C^+ a_1(t)b_1(t) + k_C^- c(t) + 2k_+ m(t)c(t) - J_1 - \sum_{i=1}^{n-1} J_i, \tag{1.4}$$

$$\frac{db_i}{dt} = J_{i-1} - J_i, \qquad 2 \le i \le n-1, \tag{1.5}$$

$$\frac{db_n}{dt} = J_{n-1}, \qquad b_i(0) = b_i^0 \quad 1 \le i \le n, \tag{1.6}$$

where $J_i$ is the net rate at which a polymer of size $3i$ grows into a polymer of size $3(i+1)$, hence:

$$J_i = k_{on}^i b_1 b_i - k_{dep}^{i+1} b_{i+1}.$$

The parameters to estimate are: $m^0$, $c^0$, $a_1^0$, $b_i^0$, $k_+$, $k_C^+$, $k_C^-$, $k_{on}^i$ and $k_{dep}^{i+1}$ - total of $2(n-1)+n+6 = 3n+4$ parameters if there are $n$ different sizes of polymers.

We can however use the properties of the model to determine part of the parameters.

#### 1.2 Interpretation of the S.V data

In the absence of an exactly reliable relation between the fraction number and the sizes of the oligomers, we assume (a choice which is qualitatively acceptable rather than exactly quantitative) that if $O_j(t)$ denotes the measured proportion of the fraction number $j$ at time $t$, it measures the proportion of polymerised mass present in oligomers containing roughly $j-$ monomers.

In the following, we denote the total polymerised mass as

$$\mathcal{M}(t) := 2a_1(t) + 3b_1(t) + 5c(t) + 3\sum_{i=2}^{n} ib_i(t).$$

We thus interpret the fraction number measured as follows:

$$\frac{2a_1(t) + 3b_1(t) + 5c(t)}{\mathcal{M}(t)} \approx \sum_{j=1}^{5} O_j(t) := \mathcal{O}_1(t), \qquad \frac{3ib_i(t)}{\mathcal{M}(t)} \approx \sum_{j=3i}^{3i+2} O_j(t) := \mathcal{O}_i(t), \quad i \geq 2,$$

and we use in the following the quantities $\mathcal{O}_i(t)$, measured at several time points, to compare the model with the experimental data. We have a maximal fraction number equal to 30, so that we define $\mathcal{O}_j$ for $j \leq 9$ and add the value of $O_{30}$ to compute $\mathcal{O}_9$ in the above definition.

Let us recall here that the size of three monomers for suPrP-B constitutes itself an approximation, so that the fit of our model to the experimental data is meant as a qualitative rather than quantitative insight.

### 2 Analysis and calibration of the model

#### 2.1 Conserved quantities

The system has two conserved quantities: first, the total mass:

$$\frac{d}{dt}\left(m + 2a_1 + 5c + 3\sum_{i=1}^{n} ib_i\right) = 0 = \frac{d}{dt}\left(m(t) + \mathcal{M}(t)\right),$$

and second, what can be viewed as the excess of monomers which will not be consumed to form suPrP-B:

$$\frac{d}{dt}\left(m - a_1 - c\right) = 0.$$

We denote these conserved quantities respectively $\mathcal{M}_{tot} = m^0 + 2a_1^0 + 5c^0 + 3\sum\limits_{i=1}^{n} ib_i^0$ and $\rho^0 = m^0 - a_1^0 - c^0$. These two quantities depend on the parameters to be estimated.

A quantity directly measured experimentally is the so-called *centroid*, defined as the average size:

$$centroid(t) := \frac{\mathcal{M}_2(t)}{\mathcal{M}(t)} = \frac{4a_1(t) + 9b_1(t) + 9\sum\limits_{i=1}^{9} i^2 b_i(t)}{2a_1(t) + 3b_1(t) + 3\sum\limits_{i=1}^{9} ib_i(t)} = \frac{\sum\limits_{j=1}^{30} jO_j(t)}{\sum\limits_{j=1}^{30} O_j(t)}.$$

#### 2.2 Asymptotic and initial behaviour of the model

We consider that at the final time measurement, an equilibrium has been reached, that we denote with $^{\infty}$ superscripts. The equilibrium fulfills the following equations:

$$\left\{ \begin{array}{rcl} -k_+ m^{\infty} c^{\infty} & = & 0, \\ k_C^+ a_1^{\infty} b_1^{\infty} - k_C^- c^{\infty} & = & 0, \\ J_1^{\infty} = \ldots = J_9^{\infty} & = & 0. \end{array} \right. \tag{2.1}$$

Asymptotically, if the monomers are in excess, the system converges towards the following state:

$$c^{\infty} = a_1^{\infty} = 0, \qquad m^{\infty} = \rho^0, \qquad b_i^{\infty} = \frac{k_{on}^{i-1}}{k_{dep}^{i}} b_1^{\infty} b_{i-1}^{\infty}, \quad i \geq 2.$$

The last equality allows us to define recursively $b_i$ from $b_1$, and $b_1$ is given by the following mass equality:

$$\mathcal{M}^{\infty} = 3 \sum_{i=1}^{9} i b_i^{\infty}.$$

Since $\mathcal{O}_1^{\infty} = \frac{3b_1^{\infty}}{\mathcal{M}^{\infty}}$, we have

$$\frac{k_{on}^{i-1}}{k_{dep}^{i}} \mathcal{M}^{\infty} = \frac{3b_i^{\infty}}{b_{i-1}^{\infty} \mathcal{O}_1^{\infty}} = 3 \frac{\mathcal{O}_i^{\infty}}{\mathcal{O}_{i-1}^{\infty} \mathcal{O}_1^{\infty}} \frac{i-1}{i}$$

which can be experimentally measured: this gives us $n-1$ relations, thus we now have $2n+5$ parameters to estimate (here $n = 9$).

We also assume that initially, before adding monomers, the system was in equilibrium, which means:

$$k_C^- c^0 = k_C^+ a_1^0 b_1^0, \qquad \frac{b_i^0}{b_{i-1}^0 b_1^0} = \frac{k_{on}^{i-1}}{k_{dep}^{i}} = \frac{\mathcal{O}_i^0}{b_1^0 \mathcal{O}_{i-1}^0} \frac{i-1}{i}, \qquad i \geq 3,$$

so that we have $n-1$ new relations, and the number of parameters to estimate is reduced to $n+6$.

#### 2.3 Numerical simulations

We run the simulations with Matlab, and used the ode solver ode45. The parameters which were not given by the analysis have been adjusted qualitatively. The time scale is in hours, the concentrations are in arbitrary units.

| | | | | | | | |
|---|---|---|---|---|---|---|---|
| $k_{on}^1$ | $k_{on}^2$ | $k_{on}^3$ | $k_{on}^4$ | $k_{on}^5$ | $k_{on}^6$ | $k_{on}^7$ | $k_{on}^8$ |
| 8.4 | 52.6 | 166 | 4.4 | 0.35 | 5.4 | 5.6 | 4.8 |
| $k_{dep}^2$ | $k_{dep}^3$ | $k_{dep}^4$ | $k_{dep}^5$ | $k_{dep}^6$ | $k_{dep}^7$ | $k_{dep}^8$ | $k_{dep}^9$ |
| 100 | 100 | 100 | 25 | 25 | 25 | 25 | 25 |
| $b_2(0)$ | $b_3(0)$ | $b_4(0)$ | $b_5(0)$ | $b_6(0)$ | $b_7(0)$ | $b_8(0)$ | $b_9(0)$ |
| 0.42 | 0.32 | 0.65 | 0.27 | 0.1 | 0.07 | 0.046 | 0.03 |
| $k_+^C$ | $k_-^C$ | $k_+$ | $m(0)$ | $c(0)$ | $a_1(0)$ | $b_1(0)$ | |
| 0.15 | 3000 | 5 | 300 | $7.10^{-4}$ | 38.2 | 0.39 | |
